# LAG3 marks activated but hyporesponsive NK cells

**DOI:** 10.1101/2024.12.18.629184

**Authors:** Valeria Vasilyeva, Olivia Makinson, Cynthia Chan, Maria Park, Colin O’Dwyer, Ayad Ali, Abrar Ul Haq Khan, Christiano Tanese de Souza, Mohamed S. Hasim, Sara Asif, Reem Kurdieh, John Abou-Hamad, Edward Yakubovich, Jonathan Hodgins, Paul Al Haddad, Giuseppe Pietropaolo, Julija Mazej, Hobin Seo, Quitong Huang, Sarah Nersesian, Damien Chay, Nicolas Jacquelot, David Cook, Seung-Hwan Lee, Giuseppe Sciumè, Stephen Waggoner, Michele Ardolino, Marie Marotel

## Abstract

Natural Killer (NK) cells are critical for immunosurveillance yet become dysfunctional in contexts such as chronic stimulation by viral infections or cancer. This phenomenon is similar to T cell exhaustion but less well characterized, which limits therapeutic interventions. As shown for T cells, NK cells often display an increased expression of immune checkpoint proteins (ICP) following chronic stimulation, and ICP blockade therapies are currently being explored for several cancer types, which have remarkable patient benefits. Nevertheless, the nature of ICP expression in NK cells is still poorly documented. In this study, we aimed to identify the conditions that lead to and the phenotype of immune checkpoint LAG3 (Lymphocyte-activation gene 3) expressing NK cells. Using various experimental models, we found that LAG3 is expressed by murine NK cells upon activation in different contexts, including in response to cancer and acute viral infections. LAG3 marks a subset of immature, proliferating and activated cells, which, despite activation, have a reduced capacity to respond to a broad range of stimuli. Further characterization also revealed that LAG3+ NK cells exhibit a transcriptional signature similar to that of exhausted CD8+ T cells. Taken together, our results support the use of LAG3 as a marker of dysfunctional NK cells across diverse chronic and acute inflammatory conditions.

## Introduction

Natural Killer (NK) cells play a role in antiviral and antitumoral immunity by directly killing stressed cells and producing immunomodulatory cytokines. Although NK cells participate in the immune response against solid and hematopoietic cancers (1), tumors often escape NK cell control. This escape is associated with a measurable loss of functional activity of NK in many types of cancers in both mice and humans (2–8). A similar hyporesponsive state of NK cells is also observed in chronic viral infections (9–13). Thus, persistent stimulation of NK cells can disarm these effector cells. Compared to T cells (14), little is known about the dysfunction of chronically stimulated NK cells, a gap in knowledge that precludes development of effective therapeutic interventions.

In the last decades, multiple efforts have been made to decipher the mechanisms governing NK cell dysfunction. These efforts were hampered by the lack of reliable surface markers to identify bona fide dysfunctional NK cells. This contrasts with the progress made in describing T cell exhaustion, whereby several molecular flags, including immune checkpoint proteins (ICP), can be used to paint a detailed molecular picture (14). To this end, we, along with other groups, have investigated whether the expression of ICPs could be used to identify dysfunctional NK cells, but obtained mixed results. For example, while programmed cell-death 1 (PD-1) is present on the surface of activated NK cells, these PD-1+ NK cells exhibit a more activated and functional phenotype than their PD-1-negative counterparts (15).

More recently, we turned our attention to lymphocyte activation gene 3 (LAG3), whose expression on exhausted T cells has been well characterized (16). LAG3 expression on NK cells was first described in the 1990s, but the function of LAG3 on these cells remains unclear. An early study in germline knockout mice revealed that LAG3-deficient NK cells exhibit reduced cytotoxic function *in vitro* (17). In contrast, blockade of LAG3 on human NK cells did not affect their ability to kill tumor cells (18). Correlative studies showed that better HIV control associated with lower expression of LAG3, PD-1, TIM-3 and CD69 on circulating NK and CD4+ T cells (19). LAG3 is also highly expressed on tumor infiltrating NK cells in patients with stage IV melanoma (20). While the combination of anti-LAG3 and anti-PD-1 therapy enhanced the cytotoxic potential of NK cells, LAG3 expression correlated with a better degranulation response against K562 target cells. Finally, LAG3 was recently proposed to act as a negative regulator of cytokine production in human NK cells (21), reviving the debate on the contribution of LAG3 to NK cell biology.

In this study, we set to investigate in which contexts LAG3 is expressed by NK cells and the phenotype associated with its expression. Using multiple models (induced and spontaneous tumors, MCMV infection, poly (I:C) or cytokine treatment), we defined common phenotypic and functional characteristics of LAG3+ NK cells. Our results revealed that LAG3+ NK cells have functional and molecular exhaustion features. Indeed, LAG3+ NK cells were more immature, activated and unresponsive to stimulation, with a transcriptional profile resembling that of exhausted CD8+ T cells. Altogether, our findings support using LAG3 as a marker of dysfunctional NK cells in studies focusing on deciphering the mechanisms that govern NK cell dysfunction.

## Results

### LAG3 is expressed on NK cells in several inflammatory contexts

To determine which stimuli trigger LAG3 expression on NK cells, we isolated murine splenic NK cells and stimulated them for 48hrs with various cytokines (IL-2, IL-15, IL-12/18 and type I IFNs). NK cells stimulated with these pro-inflammatory mediators robustly upregulated LAG3 expression (Fig. 1A, left), while PD-1 was not induced (Fig. 1A, right), consistent with the notion that murine NK cells fail to express PD-1 (15, 22). Epigenetic analysis of the *Lag3* locus revealed histone marks for promoter accessibility, such as H3K4me3, and for active enhancers, including H3K27ac and H3K4me1, that were enriched after cytokine treatment compared to that observed in untreated (NT) NK cells (Fig. 1B). Moreover, the recruitment of the acetyltransferase p300 at the LAG3 locus corroborates the increased enhancer activity induced by stimulation with IL-2+IL-12 (Fig. 1B). We next analyzed whether inflammatory cytokines induce the expression of LAG3 on NK cells in vivo, and whether this effect is cell-type specific. We treated C57BL/6 mice with poly(I:C), a Toll-like receptor-3 (TLR3) agonist often used to induce NK cell activation in vivo (23). NK cells rapidly upregulated LAG3 expression, which peaked at day 2 after stimulation (Fig. 1C, left and Fig. 1D). However, this upregulation was transient and LAG3 levels returned to baseline by 4 days post-stimulation (Fig. 1C, left). By contrast, T cells failed to upregulate LAG3 at any time point analyzed (Fig. 1C, right). Notably, LAG3 upregulation on NK cells following poly (I:C) injection was also observed in NK cells isolated from lymph nodes, the liver and the lungs (Fig. 1E). On the other hand, NK cells in the Peyer’s patches and in the small intestine, which abundantly expressed LAG3 at baseline, did not exhibit further LAG3 upregulation.

**Figure 1:**
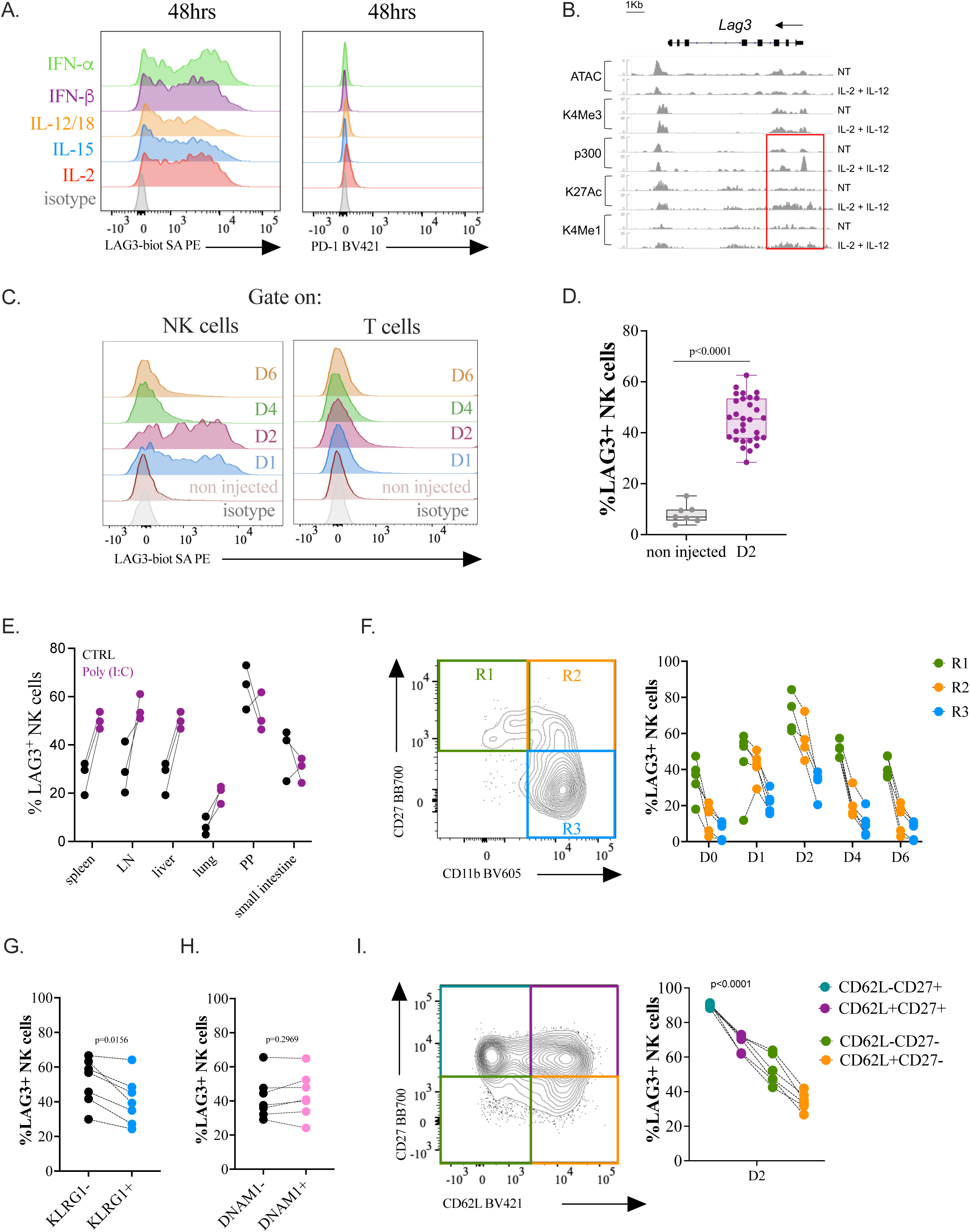
LAG3 is expressed on NK cells in several inflammatory contexts. (A) Isolated murine NK cells from C57BL/6 mice (n=4) were stimulated for 48h with different cytokines as indicated, and the expression of LAG3 and PD-1 was analyzed by flow cytometry. A representative flow plot is shown. (B) Genomic snapshots depicting chromatin accessibility (ATAC-seq), histone marks (H3K4me3, H3K27ac, H3K4me1) and p300 distribution in resting and activated NK cells *Lag3* locus. (C-D) C57BL/6 mice were injected intraperitoneally with 200ug of Poly (I:C) and LAG3 expression was analyzed by flow cytometry on splenic NK cells at various time points. (C) Representative LAG3 staining is shown on NK and T cells at various time points, n=5. (D) Mean LAG3 expression on NK cells on day 2 post injection (n=7 controls; n=30 Poly (I:C)). Data were analyzed using an unpaired non-parametric (Mann Whitney) test. (E) C57BL/6 mice were injected intraperitoneally with 200ug of Poly (I:C) and LAG3 expression was analyzed by flow cytometry on NK cells from the different organs indicated on the graph (n=3). Due to the small sample size, we did not calculate p-values. (F-I) C57BL/6 mice were injected intraperitoneally with 200ug of Poly (I:C) and LAG3 expression among CD27+CD11b-(R1), CD27+CD11b+ (R2) and CD27-CD11b+ (R3) was analyzed over time (F, n=4). Data were analyzed using a non-parametric paired Friedman test for each time point. P-value were <0.0001 except at D1, most likely because of the outlier. The expression of LAG3 among KLRG1- and KLRG1+ (G; n=6), DNAM1- and DNAM1+ (H, n=6). Data were analyzed using a paired non-parametric Wilcoxon test. The expression of LAG3 among different subsets characterized by their level of expression of CD27 and CD62L (I, n=5) was also analyzed at Day 2. Data were analyzed using a non-parametric paired Friedman test. p-values are indicated on each graph.

Next, we sought to define common phenotypic features associated with LAG3 expression. Murine CD49b+ NK cells undergo three maturation stages determined by the expression of CD27 and CD11b: immature NK cells (R1) which are defined as CD27^high^CD11b^low^, CD27 and CD11b double positive (R2) and CD27^low^CD11b^high^ mature NK cells (R3) (24). While all these NK subsets expressed LAG3 upon poly(I:C) treatment, the highest frequency of LAG3+ NK cells was found in the R1 subset, where a distinct LAG3+ population was observed not only at baseline but also after stimulation (Fig. 1F).

To confirm the higher expression of LAG3 on immature NK cells, we analyzed its co-expression with KLRG1, an inhibitory receptor that mainly marks mature and functional NK cells (25). KLRG1-NK cells expressed higher levels of LAG3 than more mature KLRG1+ counterparts (Fig. 1G). We then stratified NK cells based on the expression of DNAM-1, an activating receptor which marks an alternative functional maturation program (26). DNAM-1- and DNAM1+ NK cells expressed similar levels of LAG3 (Fig. 1H). Finally, the frequencies of LAG3+ cells were higher in a type one innate lymphoid cell like (ILC-1)-like NK cell subset (characterized by high expression of CD27 but low expression of CD62L) compared to conventional NK cells that expressed CD62L and lower levels of CD27 (27) (Fig. 1I). Collectively, these data show that NK cells upregulate LAG3 expression upon activation and that LAG3 expression is higher in less mature subsets of NK cells.

### LAG3+ NK cells are more activated and metabolically active

To elucidate potential functional differences between LAG3+ and LAG3-NK cells, we next investigated their transcriptional profile by RNA-Sequencing. LAG3+ and LAG3-murine NK cells were FACS sorted from the spleen two days after poly (I:C) injections (Fig. 2A, sorting gating strategy shown in Fig. S1A). At this time point, LAG3 was expressed on ∼50% of NK cells (Fig. 1C and S1B). In order to maximize the biological representation within each replicate, while also facilitating higher RNA yield, we pooled NK cells sorted from 10 murine spleens in each of the three replicates. The principal component analysis (PCA) revealed a clear separation between LAG3- and LAG3+ (Fig. 2B).

**Figure 2:**
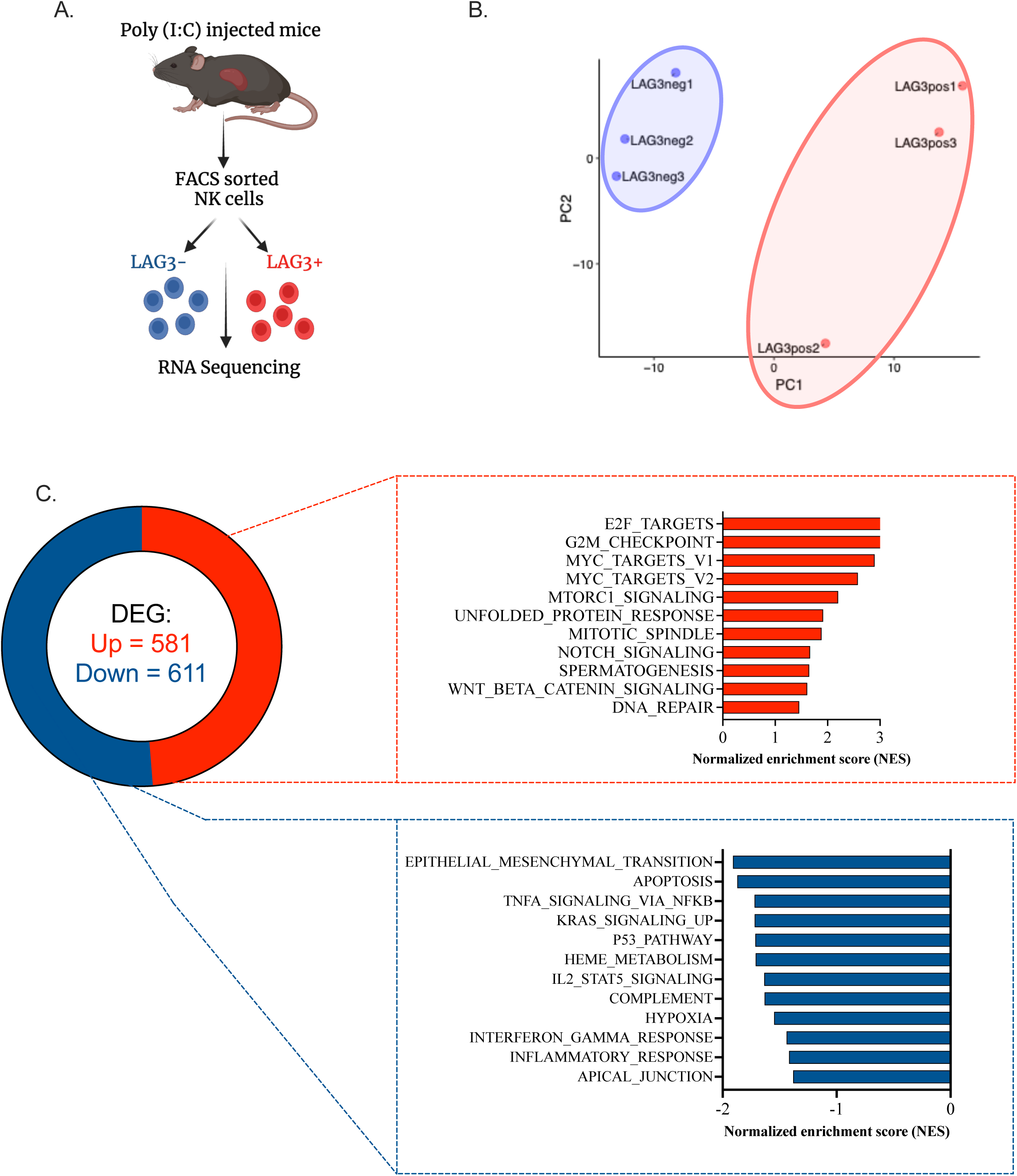
LAG3+ NK cells transcriptome suggest they are metabolically active. (A) Schematic representation of the RNA sequencing experiment. 30 C57BL/6 mice were injected with Poly (I:C). Two days post injection, mice were sacrificed and LAG3-CD11b+ and LAG3+CD11b+ splenic NK cells were sorted for RNA sequencing. (B) Principal component analysis of the RNAseq data is shown. (C) Hallmarks pathways GSEA of differentially expressed genes that are up-regulated or down-regulated in LAG3+ NK cells. Selected terms are shown among the most significant ones.

Examination of the differentially expressed genes identified 581 upregulated in LAG3+ NK cells relative to their LAG3-negative counterparts while 611 were downregulated (Log_2_ (Fold Change) >0.5 and FDR <0.05) (Fig. 2C). We performed a gene set enrichment analysis (GSEA) using the hallmark gene sets to highlight differentially active pathways in NK cells that expressed or not LAG3 (Fig. 2C). Most of the terms associating with genes enriched in LAG3+ NK cells referred to proliferation: “E2F targets, “G2M checkpoints”, “Myc target V1 and V2”, and “Mitotic spindle”. One of the other enriched terms, “mTORC1 signalling” is also associated with NK cell growth and proliferation (28). The downregulated genes in LAG3+ NK cells revealed an enrichment in the terms “apoptosis” and “p53 pathway” (Fig. 2C). Interestingly, our analysis also indicated a downregulation in pathways related to “TNFa signaling”, “IL-2-STAT5 signaling”, “IFN_Gamma response” and “Inflammatory response” collectively suggesting a reduction of genes involved in cytokine signaling. Altogether, this RNA sequencing-based profiling showed that LAG3+ NK cells exhibit a signature that seems to be characteristic of activated and proliferating cells.

### LAG3+ NK cells display higher metabolic activity and proliferative capacity

To validate the transcriptomic results, we first determined if activated NK cells presented higher expression of LAG3. Sca-1 and CD69 are commonly used as markers of NK cell activation (29, 30), hence we co-stained these two molecules with LAG3 on NK cells from mice injected with poly(I:C) (Fig. 3A). Both CD69 and Sca-1 were rapidly upregulated on splenic NK cells upon poly(I:C) injection (Fig. 3A, day 1). While CD69 expression rapidly declined at day 2, Sca-1 was more stably expressed on NK cells but approached steady state expression levels by day 4 (Fig. 3A, day 2-4). In accordance with our hypothesis, LAG3 induction on NK cells mirrored that of Sca-1, suggesting that while the cytokine response caused by poly(I:C) treatment kept NK cells activated, LAG3 was also expressed (Fig. 3A). Furthermore, Sca-1+ NK cells presented a more robust upregulation of LAG3 expression (Fig. 3B), thus confirming that LAG3 is more highly expressed by activated NK cells.

**Figure 3:**
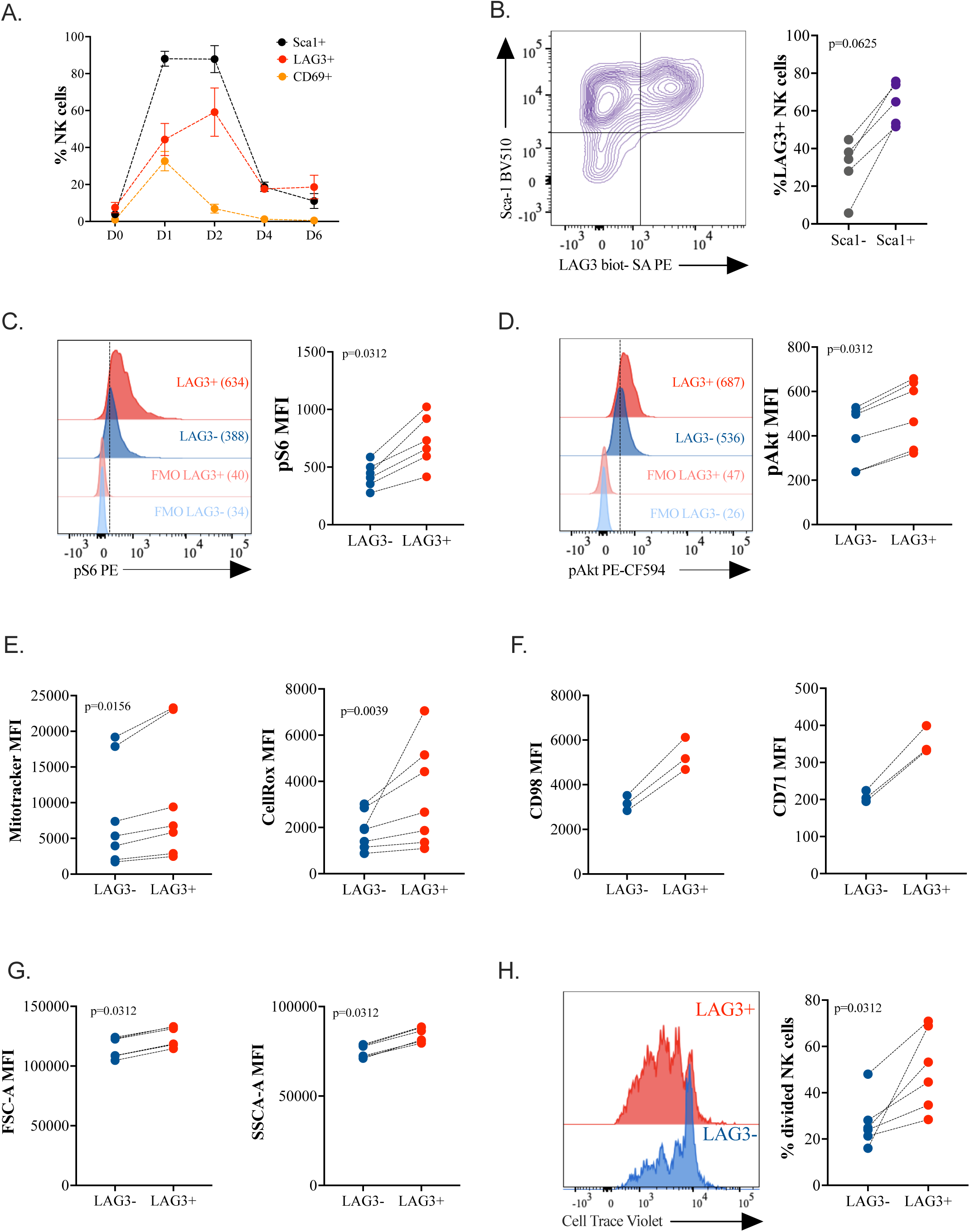
LAG3 expression indicates higher metabolic activity and proliferative capacity of NK cells. (A) C57BL/6 mice (n=4 for each day) were injected intraperitoneally with 200ug of Poly (I:C) and sacrificed in kinetic. The percentage of Sca1+, LAG3+ and CD69+ splenic NK cells was analyzed by flow cytometry. (B) The percentage of LAG3+ among Sca1+ and Sca1-NK cells on Day2 was analyzed by flow cytometry (n=5). (C-G) C57BL/6 mice (n=2 for each experiment) were injected intraperitoneally with 200ug of Poly (I:C) and sacrificed on Day2. Splenic NK cells were stained for the phospho-epitope pS6 Ser235/236 (C) and for pAkt S473 (D). Overlays of representative histograms are shown (left panels). The MFI values are indicated on the graph and represented on the right panel for each individual, n=6, 3 experiments in total. (E) NK cells were stained with Mitotracker Green and CellRox. The MFI values for each mouse (n=7, 3 experiments in total) for the analyzed marker are represented. (F) The MFI of CD98 (left) or CD71 (right) was determined by flow cytometry on LAG3+ and LAG3-NK cells (n=3). Due to the small sample size, we did not calculate p-values. (G) SSC-A and FSC-A parameters on NK cells were measured by flow cytometry in LAG3- and LAG3+ NK cells (n=5). (H) Splenic NK cells from C57BL/6 mice (n=6, 3 experiments) were isolated, stained with Cell Trace Violet (CTV) and cultured with 100ng/mL of IL-15 for 3 days. The MFI of CTV was analyzed by flow cytometry and the overlays of representative histograms are shown (left panel). The percentage of divided NK cells for each individual was calculated using the proliferation modeling tool on the FlowJo software. All the data were analyzed using a paired non-parametric Wilcoxon test. p-values are indicated on each graph.

A second hallmark pathway found upregulated in LAG3+ NK cells was mTORC1 signaling (Fig. 2C), that is required for NK cell functionality and reactivity (31, 32). Hence, we measured the basal phosphorylation level of two proteins that act downstream mTORC1/2, the ribosomal protein S6 and the kinase Akt (33) in LAG3+ vs LAG3-NK cells. LAG3+ NK cells exhibited higher phosphorylation of both S6 and Akt (Fig. 3C and 3D), indicating a stronger activity of the mTOR pathway. As mTOR plays a central role in NK cell metabolism (31), we examined the metabolic activity of LAG3+ NK cells by using two molecular probes, Mitotracker and CellRox, to respectively evaluate the global mitochondrial mass and the oxidative stress in NK cells. LAG3+ NK cells showed a significant increase in these markers (Fig. 3E), suggesting LAG3 expression tracks with metabolic activity. We then analyzed the expression of other commonly used metabolic markers such as the heavy chain of the system L amino acid transporter (CD98) and the transferrin receptor (CD71) that are both regulated by the mTOR pathway (34). LAG3+ NK cells expressed higher levels of CD98 and CD71 supporting their higher metabolic demand (Fig. 3F). Forward scatter (FSC) and Side scatter (SSC) measurements were consistently greater for LAG3+ NK cells compared to LAG3-negative counterparts (Fig. 3G), consistent with increased cell size and granularity that are additional indicators of metabolic activity in NK cells (34). Taken together, all these results suggest that LAG3+ NK cells have a higher metabolic activity and demand.

Finally, the transcriptomic analysis indicated that LAG3+ NK cells have higher proliferative capacities (Fig. 2C). To further support this observation, we analyzed the capacity of LAG3+ or LAG3-NK cells to proliferate in response to cytokines. Splenic NK cells labeled with Cell Trace Violet (CTV) were treated with IL-15 for 72 hours before assessing cell division via dilution of CTV intensity by flow cytometry. A greater proportion of LAG3+ NK cells than LAG3-NK cells diluted CTV signal (Fig. 3H), indicative of increased proliferation of NK cells expressing LAG3. These findings validate the transcriptomic analysis and confirm that LAG3+ NK cells are more activated and have higher metabolic activity and proliferative capacity.

### LAG3+ NK cells are hyporesponsive and exhibit signatures of functional exhaustion

In T cells, LAG3 and other immune checkpoint receptors are upregulated upon activation and are associated with an exhaustion program mainly driven by the transcription factor TOX (35). We thus conducted a GSEA using an independently published dataset (35) referring to the TOX-driven CD8+ T cell exhaustion signature to examine whether LAG3+ NK cells resembled exhausted T cells. The transcriptional signature of the “TOX driven exhaustion” within intra-tumoral CD8+ T cells was strongly enriched in LAG3+ NK cells (Fig. 4A). In addition to TOX, the transcriptional factors T cell factor 1 (TCF-1) and early growth response 2 (EGR2) are involved in CD8+ T cell exhaustion and expression of ICP (14, 35). Correspondingly, LAG3+ NK cells expressed more TOX, TCF-1, and EGR2 proteins (Fig. 4B).

**Figure 4:**
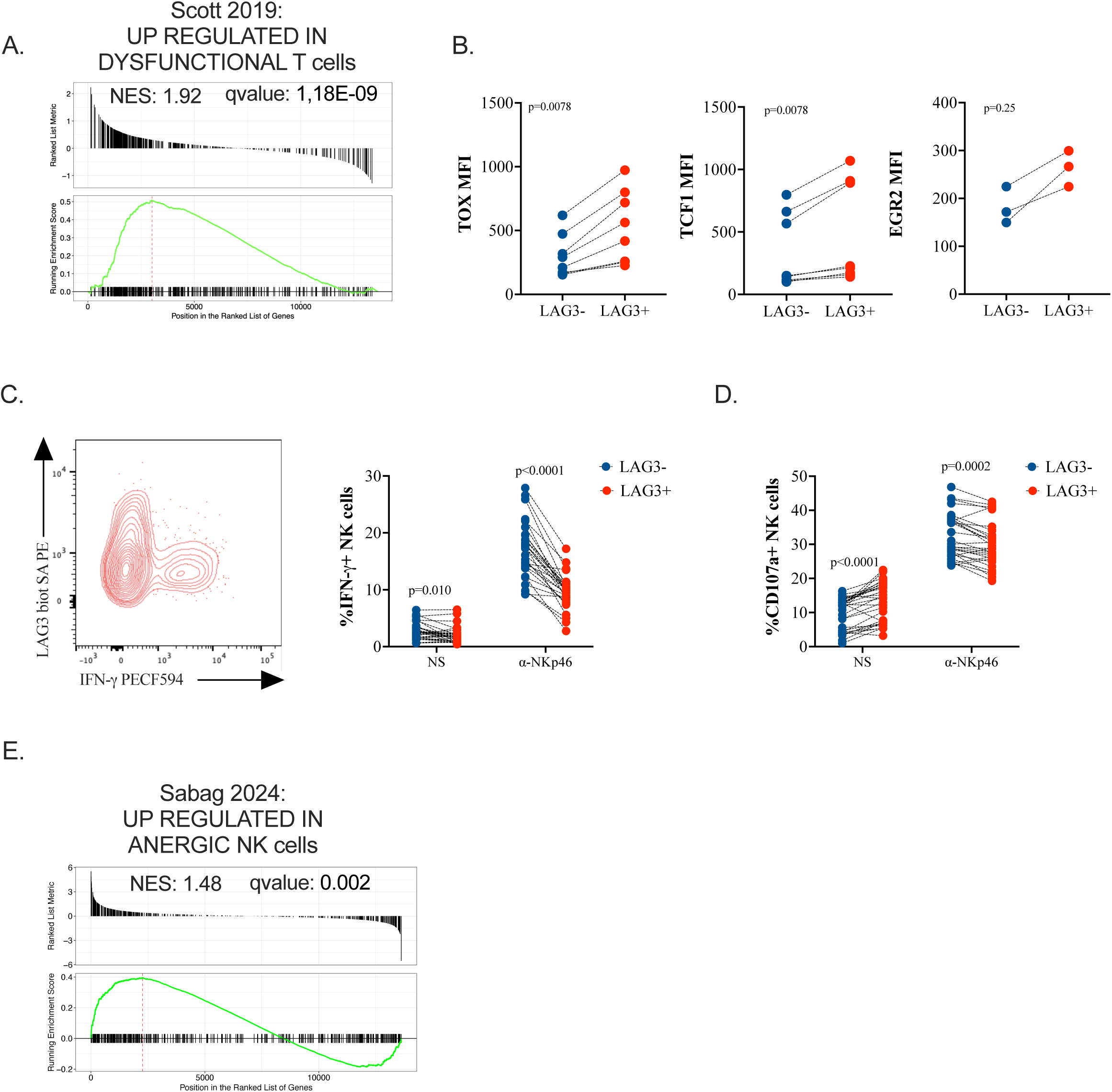
LAG3+ NK cells exhibit an exhaustion signature and are hyporesponsive. (A) GSEA plot comparing LAG3+ and dysfunctional T cells is shown for the indicated gene set. The normalized enrichment scores (NES) and FDR q-values are indicated. (B) C57BL/6 mice were injected intraperitoneally with 200ug of Poly (I:C) and sacrificed on Day 2 (n=3). The MFI values for the transcription factors TOX, TCF1 and Egr2 were determined by flow cytometry on LAG3+ vs LAG3-splenic NK cells. (C-D) C57BL/6 mice (n=30) were injected intraperitoneally with 200ug of Poly (I:C) and sacrificed at Day 2. Splenocytes were then stimulated for 4hours with plate bound NKp46 antibody. (C) Intracellular staining for IFN-γ was performed. Representative flow-cytometry plots (left) as well as proportion of IFN-γ + NK cells are shown for each individual. (D) the proportion of NK cells expressing CD107a was also determined by immunostaining and shown for each mouse. All the data were analyzed using a paired non-parametric Wilcoxon test. p-values are indicated on each graph. (E) GSEA plot comparing LAG3+ and anergic NK cells is shown for the indicated gene set. The normalized enrichment scores (NES) and FDR q-values are indicated.

Our transcriptomic and phenotypic analyses indicate that LAG3+ NK cells are activated yet present a transcriptional program similar to exhausted T cells. To more directly assess the functionality of LAG3+ NK cells, we analyzed their capacity to produce IFN-γ and degranulate (surface exposed CD107a) in response to ex vivo stimulation. Upon stimulation with NKp46, LAG3+ NK cells produced less IFN-γ and exhibited decreased degranulation compared to LAG3-NK cells (Fig. 4C-D). Of note, at the basal level (i.e. non-stimulated (NS) condition), LAG3+ NK cells displayed greater surface CD107a (Fig. 4D), suggesting they may be more activated in vivo than LAG3-NK cells. The lower reactivity displayed by LAG3+ NK cells was also apparent in response to stimulation with agonist antibodies specific for NKR-P1C or the cytokines IL-12 and IL-18 (Fig. S2A, S2B). Because LAG3 expression is highest on immature NK cells, which are known to be less responsive than mature NK cells, we stratified functional responses by maturation stage. LAG3+ NK cells exhibited reduced functional responses compared to LAG3-negative NK cells regardless of maturation stage (Fig. S2C). Taken together, these results indicate that NK cells expressing LAG3 are hyporesponsive despite their activation profile.

In addition, we observed an overlap of LAG3+ NK cells’ transcriptional profile and the signature of anergic human NK cells lacking self-MHC I inhibitory receptors as described by Sabag et al. (36) (Fig. 4E). Although the enrichment was less pronounced than the “TOX driven exhaustion” signature, the GSEA analysis revealed that key genes that are upregulated in anergic NK cells were also enriched in LAG3+ NK cells, further confirming their hyporesponsive state.

### LAG3 expression is a shared feature of NK cell activation

We next sought to ascertain whether LAG3 is a marker of NK cells with reduced functions in the context of viral infections and cancer. First, we took advantage of the murine cytomegalovirus model (MCMV), which has been extensively used to study the role of NK cells in viral infections (37, 38). C57BL/6 mice were infected with MCMV (15,000 PFU) and sacrificed at D2, D4, D7 and D10. We found that, unlike T cells, LAG3 expression was enhanced on splenic NK cells from mice infected with MCMV, in particular at D2 and D4 post-infection (Fig. 5A). This was also true for liver NK cells (Fig. S3A, left), while hepatic T cells expressed LAG3 with a delayed kinetic (Fig. S3A, right). A higher percentage of NK cells expressed LAG3 within the activated Sca1+ population in both the spleen (Fig. 5B) and the liver (Fig S3B). In contrast, similar LAG3 levels were expressed by Ly49H+ and Ly49H-NK cells (Fig. S3C and S3D), suggesting that the inflammatory response, and not antigen recognition, may be primarily driving LAG3 expression on NK cells.

**Figure 5:**
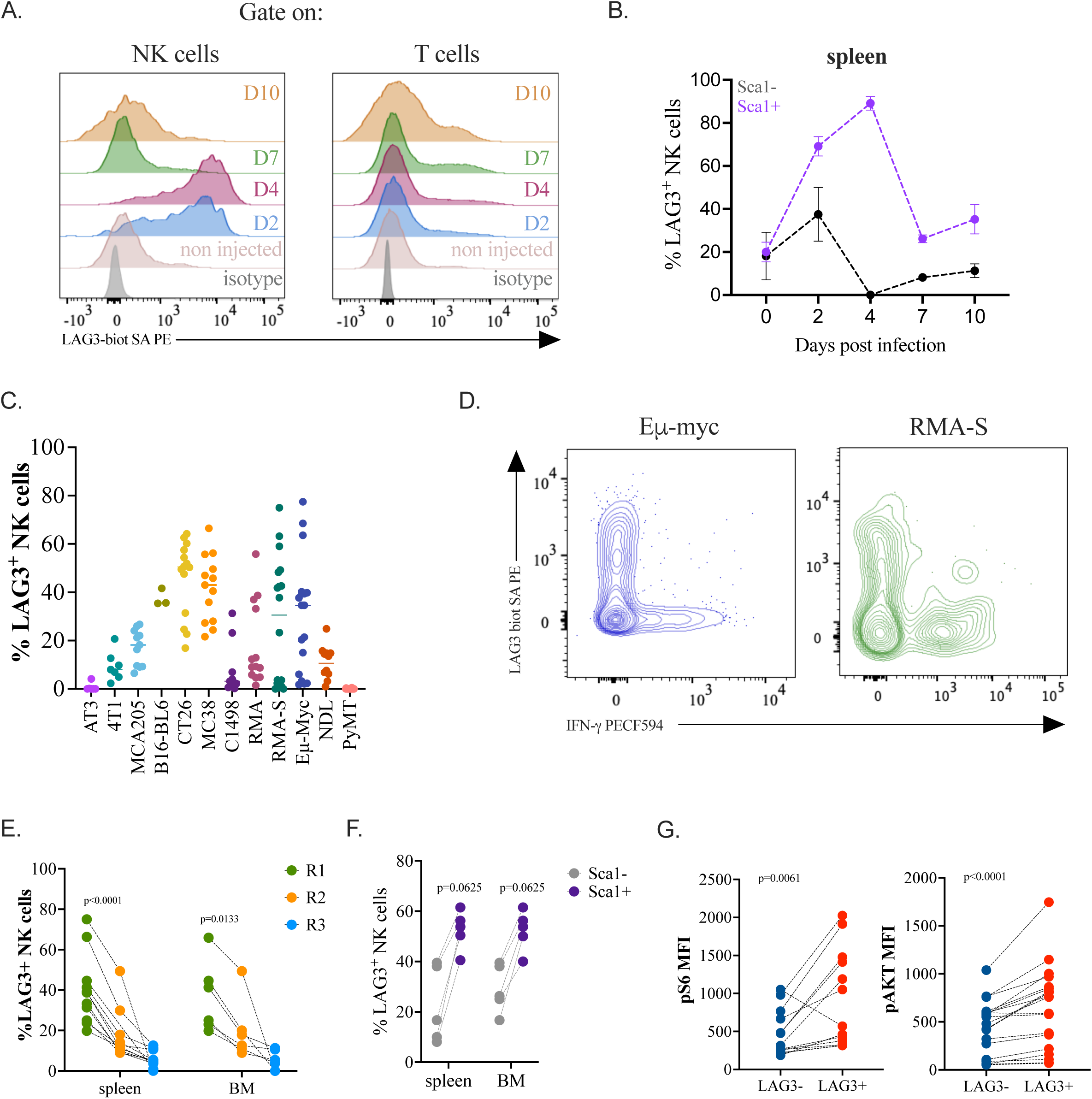
LAG3 expression is a shared feature of NK cell activation. (A-B) C57BL/6 mice (n=3) were infected with MCMV (15,000 PFU) and sacrificed in kinetic at D2, D4, D7 and D10. The expression of LAG3 on splenic NK and T cells was determined by flow cytometry. Overlays of representative histograms are shown in NK cells (A, left panels) or in T cells (B, right panels). (B) The percentage of LAG3 among Sca1+ or Sca1-splenic NK cells was determined by flow cytometry. Due to the small sample size, we did not calculate p-values. (C) Summary of LAG3 expression on intra-tumoral NK in mice injected with AT3, 4T1, MCA205, B16-BL6, C1498, RMA, or RMA-S or on intra-tumoral NK cells in the spleen or tumors from spontaneous cancer models (Eµ-Myc models or NDL and PyMT, respectively). LAG3 expression on NK cells in each model was assessed in at least 3 independent experiments with at least n = 3 except for the B16-BL6 sub-cutaneous model. (D) Splenocytes from spontaneous cancer models (Eµ-myc, n=5)) or intra-tumoral NK cells from ectopic tumor model (RMA-S, n=3)) were stimulated for 4hours with plate bound NKp46 antibody. Intracellular staining for IFN-γ was performed. Representative flow-cytometry plot is shown (Eµ-myc, n=5 and RMA-S, n=3). (E) LAG3 expression among CD27+CD11b-(R1), CD27+CD11b+ (R2) and CD27-CD11b+ (R3) in NK cells (from the spleen or Bone Marrow) from the Eµ-myc spontaneous model (n=5) was analyzed by flow cytometry. Data were analyzed using a paired non-parametric Friedman test. (F) LAG3 expression among Sca1- or Sca1+ NK cells (from the spleen or Bone Marrow) from the Eµ-myc spontaneous model (n=5) was analyzed by flow cytometry. Data were analyzed using a paired non-parametric Wilcoxon test. (G) Splenic NK cells from the Eµ-myc spontaneous model were stained for the phospho-epitope pS6 Ser235/236 (left panel) and for pAkt S473 (right panel). The MFI values for each mouse are indicated on the graph, n=13. Data were analyzed using a paired non-parametric Wilcoxon test. p-values are indicated on each graph.

We next investigated if NK cells upregulated LAG3 in models of cancer. LAG3 was abundantly expressed on tumor-infiltrating NK cells in various ectopic or spontaneous murine tumor models (Fig. 5C). NK cells infiltrating the tumor models analyzed often presented LAG3 expression, although with some degree of heterogeneity. We found the highest expression in the colon cancer CT26 and MC38 models, in the T cell lymphoma ectopic tumor RMA-S model and in the spontaneous B cell lymphoma Eµ-Myc model.

To further characterize LAG3+ intra-tumoral NK cells, we focused on the Eµ-Myc tumor model, where mice spontaneously develop a B cell lymphoma in several organs (bone marrow, lymph nodes, spleen and thymus). This model allowed us to recover a higher number of NK cells from tumors for subsequent analyses. Importantly, LAG3+ NK cells were highly hyporesponsive in the Eµ-Myc model (Fig. 5D, left). A similar dissociation in expression of LAG3 and IFN-γ was observed in NK cells infiltrating RMA-S ectopic tumors (Fig. 5D, right). Importantly, the profile of tumor infiltrating NK cells expressing LAG3 resembled those induced by poly (I:C) in terms of enrichment of an immature (CD27+ CD11b-, KLRG1-, DNAM1-) and activated (Sca1+) phenotype (Fig. 5E-F, 2G-H) with higher mTOR signaling (pS6 and pAkt) activity (Fig. 5G). Altogether, these results show that, in tumor models, LAG3 is more expressed by immature NK cells that are activated yet hyporesponsive and suggest that LAG3 could be used as a marker of dysfunctional NK cells.

### LAG3 is not required for NK cell development nor for acute killing of RMA-S

The strong association between an activated, hyporesponsive phenotype and expression of LAG3 in NK cells prompted us to employ a genetic tool to mechanistically link LAG3 expression with NK cell dysfunction. To this end, we obtained *Lag3^fl/fl^*mice generated on a mixed C57BL/6 and 129 background (39) that were extensively backcrossed to fix critical components of the C57BL/6 NK receptor complex (NKC) located on chromosome 6. Of note, Lag3 is located close to the NKC on the same chromosome. We then crossed *Lag3^fl/fl^* mice with *Ncr1*-iCre mice to generate a mouse line lacking LAG3 expression selectively in NK cells (termed LAG3-KO) that can be compared to *Cre*-negative LAG3-WT littermates (Fig. 6A). To validate LAG3 deletion in this model, we injected mice of both genotypes with poly(I:C) and analyzed the expression of LAG3 on NK cells at Day 2. Poly(I:C) injection induced LAG3 expression exclusively on NK cells from LAG3 WT mice but not in LAG3 KO mice (Fig. 6B), with levels comparable to those observed in C57BL/6 mice treated with poly(I:C) (Fig. 1D).

**Figure 6:**
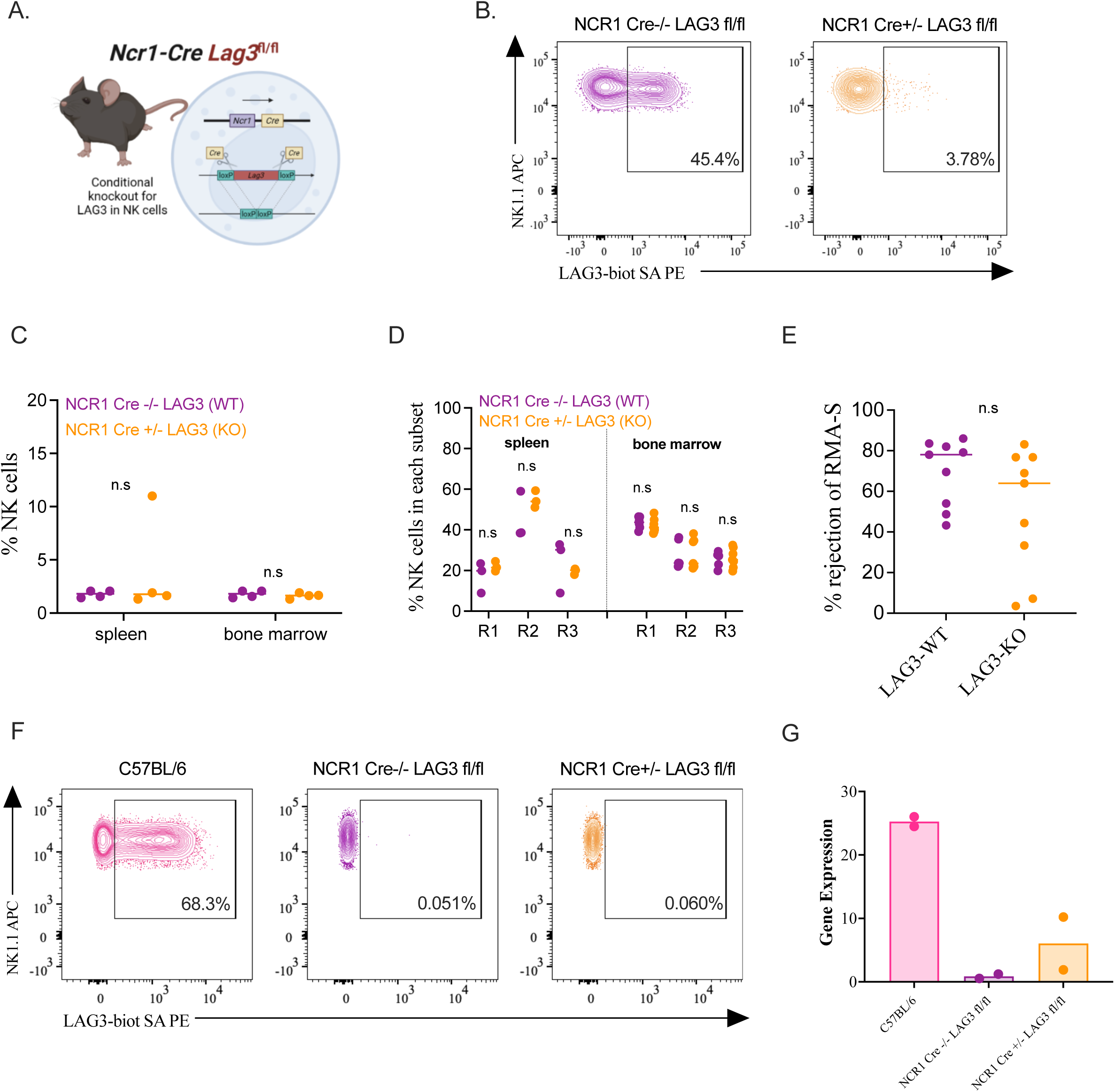
LAG3 marks a subset of hyporesponsive NK cells but may not be the driver of the dysfunctional phenotype. (A) Schematic representation of the NCR1-iCre LAG3-flox model. (B) NCR1 Cre-/- LAG3 fl/fl (LAG3-WT) and NCR1 Cre+/- LAG3 fl/fl (LAG3-KO) were injected with 200ug Poly (I:C) and sacrificed at Day2. LAG3 expression on splenic NK cells was determined by flow cytometry. Representative flow plots are depicted (out of n=5). (C-D) NCR1 Cre-/- LAG3 fl/fl (LAG3-WT) and NCR1 Cre+/- LAG3 fl/fl (LAG3-KO) were sacrificed at 8-9 weeks old (n=5 for each group). (C) The percentage of NK cells in the spleen and bone marrow was determined by flow cytometry. (D) The percentage of NK cells in each maturation subset in the spleen and bone marrow was determined by flow cytometry. (E) NCR1 Cre-/- LAG3 fl/fl (LAG3-WT, n=9)) and NCR1 Cre+/- LAG3 fl/fl (LAG3-KO, n=9) were injected with 200ug Poly (I:C). 24h later, they were injected with a 50:50 mixture of RMA/RMA-S cells. 24h later, the percentage of remaining RMA-S and RMA cells in the peritoneal wash was determined by flow cytometry. All data were analyzed using an unpaired non-parametric (Mann-Whitney) test. p-values are indicated on each graph. (F-G) C57BL/6, NCR1 Cre-/- LAG3 fl/fl (LAG3-WT) and NCR1 Cre+/- LAG3 fl/fl (LAG3-KO) were injected with 200ug Poly (I:C) and sacrificed at Day2 (n=2 per group). LAG3 expression on splenic NK cells was determined by flow cytometry (F). Representative flow plots are depicted (out of n=2 per group). (G) qPCR of LAG3 gene was performed on these NK cells.

We then assessed potential defects in the NK cell compartment due to the absence of LAG3. LAG3-KO mice presented a normal number of NK cells (NK1.1+NKp46+) in the spleen and bone marrow (Fig. 6C) and no apparent defect in phenotypic maturation of NK cells (Fig. 6D).

To assess the role of in NK cell anti-tumor activity, we performed *in vivo* killing assays in groups of mice treated with poly(I:C) to induce LAG3 expression prior to injection of a 50:50 mixture of RMA (NK cell resistant): RMA-S (NK cell sensitive) tumor cells. After 24 hrs, we analyzed the percentage of remaining RMA-S cells relative to RMA cells as an indicator of NK cell cytotoxicity. In this setting, LAG3 KO mice cleared RMA-S similarly to littermate control, indicating that LAG3 is dispensable for acute killing of MHC-deficient tumor cells (Fig. 6E).

Unfortunately, an unexpected shift in LAG3 expression occurred after these initial studies in the LAG3 mouse colonies that calls for caution in the use of this model. While LAG3 was properly upregulated in NK cells in Cre negative control mice when the colony was first established and evaluated (November 2020; Fig. 6A-E), this response waned in the control mice over time. In subsequent experiments we failed to observe an induction of LAG3 expression in Cre-negative LAG3 WT littermate control NK cells after treatment of mice with poly(I:C), in contrast to continued expression of LAG3 on NK cells in poly(I:C)-treated C57BL/6 controls (Fig. 6F). Lack of LAG3 induction at the protein level was associated with a corresponding failure to upregulate *Lag3* expression at the mRNA level (Fig. 6G). Therefore, our littermate *Ncr1*-Cre-/- *Lag3^fl/fl^* colony represents a poor control when it comes to investigating the role of LAG3 in controlling viral infections or tumor development.

## Discussion

Immune cell dysfunction is a complex and multifaceted phenomenon that remains not fully understood. Therefore, it is essential to define precise markers that can identify subsets of these dysregulated immune cell populations. In this study, we focused on identifying the receptor LAG3 as a marker for dysfunctional NK cells. LAG3 was found to be expressed by activated T and NK cells (40, 41) and its expression was later reported on B cells (42), NKT cells (43), plasmacytoid dendritic cells (44) or regulatory T cells (T_regs_) (45). LAG3 is a well-established marker of CD8+ T cell exhaustion, particularly in viral and tumor settings, with evidence showing that blocking LAG3 restores T cell function in cancer models (46–48). This highlights LAG3’s potential role in not only driving but also maintaining T cell exhaustion. In our hands, LAG3 was broadly expressed in vitro and in vivo on NK cells upon activation, whereas a lower expression was observed in T cells. Conversely, T cells expressed high levels of PD-1, which was not detected on NK cells, potentially suggesting that different immune cells may have different checkpoint receptors. Recent studies have clarified LAG3’s specific functions in modulating T cell exhaustion, including control over cytotoxicity, cytokine polyfunctionality, and NK cell receptor expression distinct from PD-1, which influences T cell expansion. LAG3 also plays a role in promoting the expression of TOX, an epigenetic regulator of exhaustion, with LAG3 deletion enhancing CD8+ T cell effector functions (49). In our hands, LAG3+ NK cells have a similar transcriptomic signature to exhausted CD8+ T cells (35). LAG3+ NK cells showed increased expression of the key drivers of T cell exhaustion: Tox, Tcf1, and Egr2 (50–52). Inhibition of Egr2 rescues NK cell dysfunction (36), but TOX and TCF1 are involved in NK cell development (53, 54) and have yet to be shown to contribute to NK cell dysfunction. In Tregs, LAG3 is essential for suppressive activity, affecting the PI3K-Akt-Rictor pathway and Myc-driven metabolic programming (55). Despite these well-defined roles in T cells and Tregs, LAG3’s function in NK cells remains less understood and warrants further investigation.

Our data suggests that NK cells expressing LAG3 are activated but hyporesponsive, a phenotype that is consistent with T cell exhaustion. Since NK cells react quickly to stimulation, the rapid expression of LAG3 seems appropriate based on their fast-acting nature (56). In our hands, other immune checkpoints like PD-1, TIGIT, and TIM3 were only modestly upregulated or absent on NK cells in inflammatory conditions and in cancer in comparison to LAG3. LAG3 expression is also observed in previous studies investigating NK cells under chronic condition suggesting that this protein may be important in NK cells facing stimulation whose fate may be akin to exhausted CD8+ T cells (13). Thus, we propose that LAG3 could serve as a reliable marker for identifying certain dysfunctional NK cell populations.

Dysfunctional immune cell populations are heterogeneous, exhibiting distinct phenotypes that vary depending on the specific context. We suggest that LAG3 can be used to identify a subset of dysfunctional NK cells that are overactivated yet hyporesponsive. This phenotype resembles other dysfunctional immune subsets such as T cells undergoing early stages of exhaustions referred to as precursors to exhausted CD8+ T cells (Tpex) (57). LAG3+ NK cells share similarities with Tpex cells, including increased expression of TCF-1 (a transcriptional factor decreased in canonically exhausted CD8+ T cells (58)), elevated proliferation, greater mitochondrial mass, and activation of the mTOR pathway. This could reflect a unique dysfunctional phenotype in NK cells, distinct from canonically exhausted CD8+ T cells, due to NK cells having shorter lifespans, a lack of antigen specificity, and a rapid immune response activation (59–61). However, studies have shown that T cell function and proliferation in response to stimulation are uncoupled (62), and overstimulation may trigger two separate exhaustion programs (58). Like T cells (63), it is proposed that NK cells expressing LAG3 might be attempting to balance effector function and prevent overstimulation by becoming hyporesponsive to simultaneously provide modest control of harmful cells without significant tissue damage (63). Other immune checkpoints like CTLA-4, TIGIT, and TIM3 also show context-dependent expression on NK cells, but it remains unclear whether they simply also mark dysfunction or drive it. Recent findings suggest that immune checkpoints, including LAG3, are expressed on dysfunctional NK cells but are not the primary cause of functional impairment, indicating that other mechanisms contribute to NK cell dysfunction (64), Taken these findings into account, we could further speculate that LAG3 may serve as a marker for identifying an NK cell population balancing aggressive immune responses with restraint to prevent immunopathology.

Further investigation into the role of LAG3 in NK cells is crucial to fully understand its significance in overly activated, hyporesponsive NK cells. However, the currently available NK cell-targeted LAG3 knock out murine model seems not suitable for further investigations since our data revealed that LAG3 expression is lost in the WT littermate controls. This loss of gene expression could be due to unpredictable genetic changes near *Lag3* on chromosome 6 arising from attempts to backcross to C57BL/6 background based on nearby NK complex (NKC) locus. In conclusion, in our study, we have identified LAG3 as a marker of NK cell dysfunction, offering a valuable tool for identifying dysfunctional NK cells in future studies and leading to a better understanding of the mechanisms underlying this phenomenon in different pathological conditions and immunotherapies. Surprisingly, we found that immune cells exhibiting reduced functionality, such as LAG3+ NK cells, are paradoxically more metabolically active, likely due to their overactivation. It will be important to follow up on these data, particularly in light of ongoing efforts to use LAG3 blocking reagents in the clinic (65).

## Material and Methods

### Animals

Animals were housed and maintained at the University of Ottawa Animal Care and Veterinary Services (ACVS) or at the University of Calgary (AC23-0054; AC23-0003; AC23-0162) with standard protocols for housing, diet, and care. Animal use was approved by the University of Ottawa Animal Ethics and Compliance Board. C57BL/6 mice, NCR1-GFP and Eµ-Myc mice were purchased from Jackson Laboratories. Both male and female 8–15-week-old were used. Eµ-Myc mice were sacrificed when they spontaneously developed B cell lymphoma and experienced cancer symptoms, such as enlarged masses on the neck and lethargy, around 11-15 weeks. NCR1-Cre LAG3-flox mice were obtained from Dr. Stephen Waggoner who initially obtained LAG3^fl/fl^ mice from Dr. Dario Vignali (University of Pittsburg)(39). NDL mice were a kind gift from Dr. Luc Sabourin (Ottawa Hospital Research Institute). PyMT mice were kindly provided by Dr. Pamela Ohashi (University of Toronto) who obtained them from Dr. William Muller (Mc Gill) (66).

### Intraperitoneal poly I:C injection

C57BL/6 mice were intraperitoneally injected with 200 µg of polyinosinic:polycytidylic acid (poly I:C) (Invivogen). Mice were sacrificed at different time points after injection.

### Tumor injections

For s.c. and orthotopic injections, tumor cells resuspended in 100 μl RPMI without FCS were injected in the left flank or in the mammary fat pad. For in vivo killing assay, a mixture composed of 5 million RMA cell and 5 million RMA-S cells were injected intra peritoneally. 24hours later, we performed peritoneal washes to collect the content of the peritoneal cavity and analyzed by flow cytometry the percentage of remaining RMA-S cells relative to RMA cells as an indicator of NK cell cytotoxicity.

### Tumor dissociation

Tumors were collected at endpoint, cut in small pieces and dissociated using the GentleMacs (Miltenyi^TM^) in media supplemented with DNase I and collagenase IV for 45 min at 37°C. Cells were then homogenized using Fisherbrand Sterile Cell Strainers and resuspended in Phosphate Buffered Saline (PBS) (1X, pH 7.4). Red blood cell lysis was performed using homemade Ammonium Chloride-Potassium (ACK) lysing buffer. Cells were resuspended in 5% Gibco^TM^ RPMI prior further analyses.

### Splenocyte isolation

Spleen was homogenized using Fisherbrand Sterile Cell Strainers and resuspended in Phosphate Buffered Saline (PBS) (1X, pH 7.4). Red blood cell lysis was performed using homemade Ammonium Chloride-Potassium (ACK) lysing buffer. Splenocytes were resuspended in 5% Gibco^TM^ RPMI prior further analyses.

### NK cell isolation

Murine NK cells were isolated from splenocytes using the Mouse NK cell Isolation Kit (StemCell Technologies EasySep^TM^) according to manufacturer’s instructions.

### In vitro stimulation assay

Isolated NK cells were cultured for 48h at 200,000 cells per mL in presence of different cytokines: 1,000 IU/mL of type I IFN-⍺, 1,000 IU/mL type I IFN-β, 20ng/mL of IL-12 + 100ng/mL of IL-18, 100 ng/mL of IL-15, or 1,000 IU/mL IL-2.

### Flow cytometry analysis

Isolated splenocytes were incubated with BD Biosciences Mouse Fc Block^TM^ Purified Rat Anti-Mouse CD16/CD32 to block FcγRII/III receptors. Cells were then immunostained during 30 min at 4°C with the appropriate monoclonal antibodies detailed list in the Supplementary Table 1. For phosphoflow assays, cells were then incubated for 1 hour in 5% Gibco^TM^ RPMI at 37℃ to preserve LAG3 extracellular staining. Intracellular staining of cytokines and chemokines was performed with Cytofix/Cytoperm (BD Biosciences). Intracellular staining of transcription factors or cytotoxic molecules was performed using Foxp3 Fixation/Permeabilization concentrate and diluent (eBioscience). Intracellular staining of phosphorylated proteins was performed with Lyse/Fix and Perm III buffers (BD Biosciences) according to the manufacturer’s instructions. Phosphorylated proteins were then stained during 40 min at RT with the appropriate antibodies detailed in the Supplementary Table 1. Stained cells were resuspended in homemade flow cytometry staining buffer (PBS + 0.5% BSA) and analyzed using flow cytometry. Flow cytometric analysis was performed on LSR Fortessa 5L (Becton-Dickinson). Fluorescence Minus One or IgG controls were used to set the gates, and data were analysed with FlowJo 10.9.0 software.

### Ex vivo functionality assay

Isolated splenocytes were collected and stimulated using the plate-bound antibodies against NK cell activating receptors, NKp46 (BD Biosciences^TM^ Purified Rat Anti-Mouse CD335/NKp46), NK1.1 (Stem Cell Technologies, PK136 clone) and NKG2D (Invitrogen^TM^ Monoclonal Antibody NKG2D) or with Recombinant Murine IL-12 (20ng/mL, Peprotech) and Recombinant Murine IL18 (100ng/mL, Peprotech) cytokines for 4 hours at 37℃ in presence of the CD107a antibody and Golgi Stop and Plug (BD biosciences) The percentage of CD107a+ and IFN-γ+ cells was analyzed by flow cytometry. Isolated splenocytes were extracellularly stained for LAG3 and exhaustion markers.

### In vitro proliferation assay

Isolated NK cells were stained with 0.6µM Invitrogen CellTrace^TM^ Violet (CTV) Cell Proliferation Kit according to manufacturer’s instructions. Stained NK cells were then incubated in 5% Gibco^TM^ RMPI with 100ng/mL PeproTech^TM^ Recombinant Murine IL-15 for 72 hours at 37℃. After 72 hours, cells were analyzed using flow cytometry. The percentage of divided NK cells for each individual was calculated using the proliferation modeling tool on the FlowJo 10.9.0 software.

### Ex vivo oxidative stress detection assay

Isolated splenocytes were stained extracellularly for NK markers and LAG3. Then cells were incubated with CellRox^TM^ Deep Red Reagent (Invitrogen) in PBS during 30 min according to manufacturer’s instructions and then assessed using flow cytometry.

### Ex vivo mitochondrial content detection assay

Isolated splenocytes were stained extracellularly for NK markers and LAG3. Then cells were incubated with MitoTracker^TM^ Deep Red Reagent (Invitrogen) in PBS during 10 min according to manufacturer’s instructions and then assessed using flow cytometry.

### Cell sorting for RNA sequencing

NK cells were sorted as CD3-CD19-NK1.1+NKp46+DX5+LAG3+ or CD3-CD19-NK1.1+NKp46+DX5+LAG3-as shown in Supplementary Figure S1, using a Beckman Coulter MoFlo XDP cytometry sorter.

### RNA Isolation for RNA sequencing

RNA from sorted NK cells was isolated using the Sigma^TM^ GenElute RNA Miniprep Kit as per manufacturer’s protocol. mRNA sequencing libraries were prepared according to the manufacturer’s instructions for the TruSeq Stranded mRNA Library Prep kit (Illumina, San Diego, California, USA). Library preparation and sequencing was performed by DNA Link (DNA Link Inc., Seoul, Republic of Korea). mRNA was purified and fragmented from total RNA using poly-T oligo-attached magnetic beads using two rounds of purification. Cleaved RNA Fragments primed with random hexamers were reverse transcribed into first strand cDNA using reverse transcriptase, random primers, dUTP in place of dTTP. These cDNA fragments then had the addition of a single ’A’ base and subsequent ligation of the adapter. The products were purified and enriched with PCR to create the final strand specific cDNA library. The quality of the libraries was verified by Tapestation (Agilent Technologies, Santa Clara, CA, USA). After quantitative PCR (qPCR) using KAPA SYBR FAST qPCR Master Mix (Kapa Biosystems, Wilmington, MA, USA), we combined libraries that index tagged in equimolar amounts in the pool. Sequencing was performed using an Illumina NovaSeq 6000 system following provided protocols for 2 × 100 sequencing. Transcript quantification for each sample was performed using Kallisto (v0.45.0) (66) with the GRCm38 transcriptome reference and the -b 50 bootstrap option. The R package DESeq2 (v1.44.0) was then used to construct general linear models of each gene across experimental conditions. Wald’s test was used to test for differential expression between groups and the resultant p-values were adjusted using the Benjamini-Hochberg false discovery rate method.

### RNA isolation and preparation for qPCR

RNA from sorted NK cells was isolated using the Sigma GenElute^TM^ RNA Miniprep Kit as per manufacturer’s protocol. Using the BioRad iScript^TM^ Reverse Transcription Supermix kit, cDNA was obtained alongside a no reverse transcriptase control and RNAase/DNAase free water control using a PCR machine. The samples were assessed for purity using a Nanodrop. Using the Biored iTaq^TM^ UniversalSYBR Green Supermix kit, forward and reverse primers, the qPCR samples were prepared using the ThermoFisher 7500 fast Real-Time PCR machine.

### qPCR analysis

Analysis of relative gene expression data using real-time quantitative PCR was done according to a study performed by Kenneth J. Livak and Thomas D. Schmittgen (Analysis of Relative Gene Expression Data Using Real-Time Quantitative PCR and the 2−ΔΔCT Method, Methods, Volume 25, Issue 4, 2001).

### Epigenetic analyses of LAG3 locus

ATAC-Seq/ChIP-Seq were previously generated and mapped onto mouse genome build mm9, as described (67). Genomic snapshot of the *Lag3* locus was generated using IGV software (version 2.8.2) (68).

### Statistical Analysis

Univariate statistical analyses were performed on GraphPad Prism. Unpaired or paired statistical test was used as appropriate and indicated in each figure legend.

## Acknowledgements

We thank members of the Ardolino laboratory and all the authors for critically reading the manuscript; The OHRI and uOttawa flow-cores for support with flow cytometry; and the ACVS facility at the University of Ottawa for support with animal studies. We are thankful to Dr. Vignali (University of Pittsburg) for providing the LAG3 fl/fl mice, Dr. Ohashi (University of Toronto) for providing the PyMT mice and Dr. Sabourin (OHRI) for providing the NDL mice.

## Author contributions

Conceptualization: M.M and M.A. Funding acquisition: MM and M.A. Investigation: V.V, M.M, O.M, C.C, M.P, C.D, A.A, C.T.D.S, M.S.H, S.A, R.K, J.A, E.Y, J.H, P.A.H, G.P, J.M, H.S, Q.H, G.S and M.A. Methodology: M.M. and M.A. Resources: G.S, S.W, D.Co, N.J, Supervision: A.S., G.S., A.Z., and M.A. Visualization: V.V, C.D, S.N, D.C, D.Co and M.M. Writing—original draft: V.V, M.M, S.N, S.W and M.A. Writing—review and editing: all authors.

---

**Figure S1:**
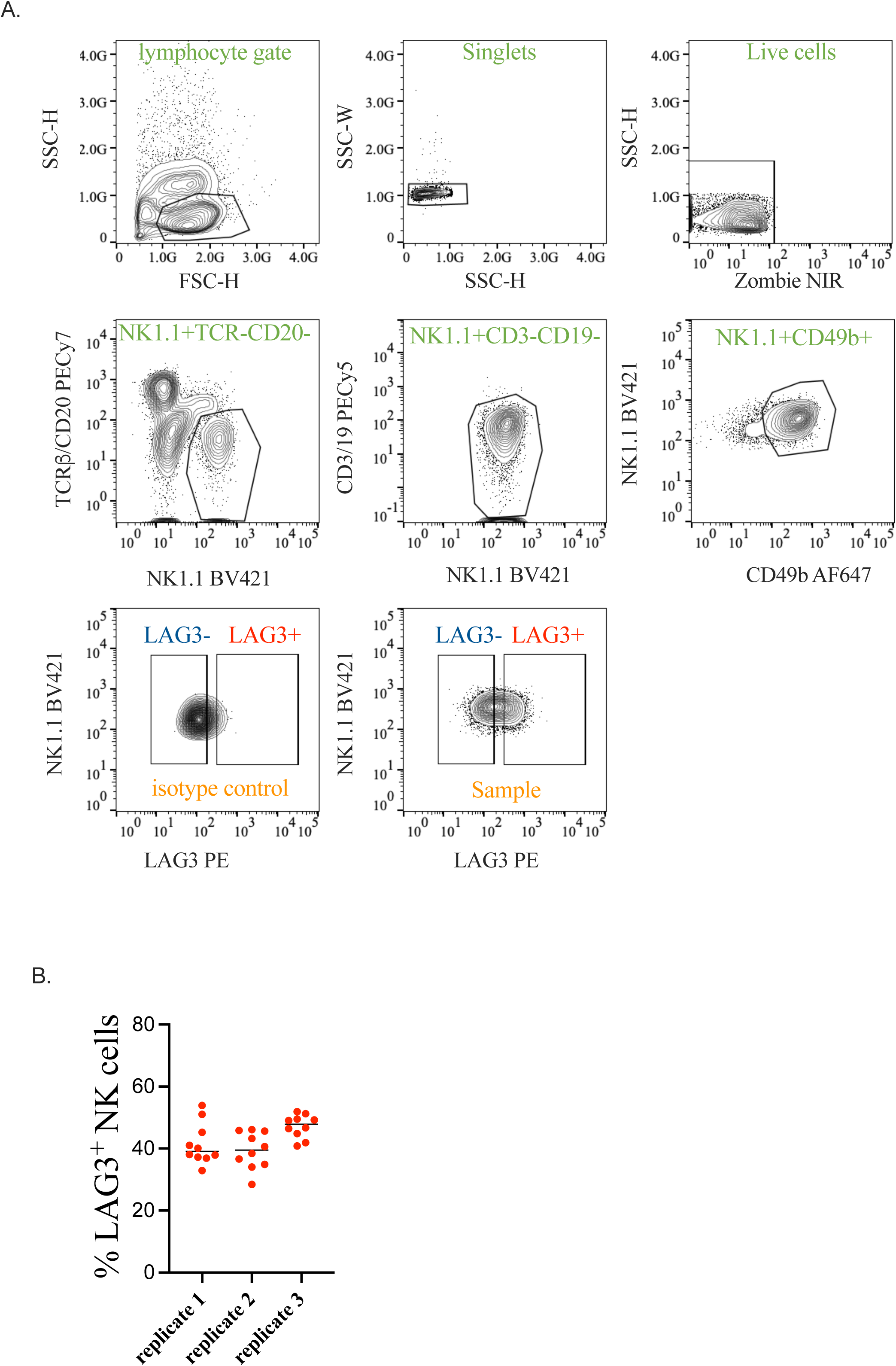
(A) Gating strategy used to sort LAG3+CD11b+ and LAG3-CD11b+ NK cells from the spleen of mice injected with poly (I:C) for RNA sequencing (B) LAG3 expression on NK cells from the spleen of mice injected with poly (I:C) was analyzed by flow cytometry. The percentage of LAG3+ NK cells in each mouse (n=30) for the three replicates used for the RNA sequencing is depicted on the figure.

**Figure S2:**
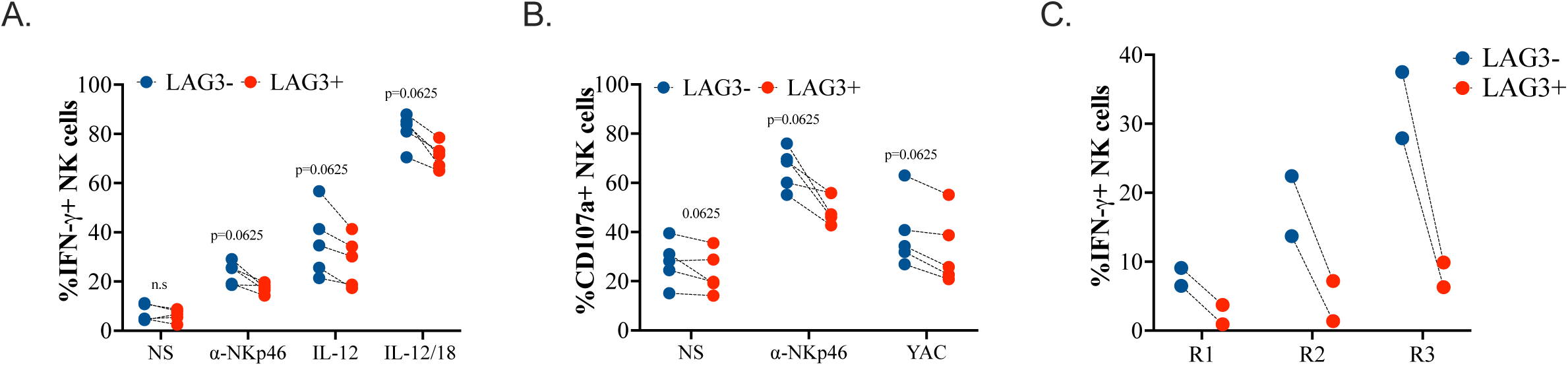
(A-B) C57BL/6 mice (n=4) were injected intraperitoneally with 200ug of Poly (I:C) and sacrificed at Day 2. Splenocytes were then stimulated for 4hours with plate bound NKp46 antibody, with IL-12, IL-12+IL-18 or with YAC-1 cells at a 1:1 ratio. (A) Intracellular staining for IFN-γ was performed. The proportion of IFN-γ + NK cells are shown for each individual. (B) the proportion of NK cells expressing CD107a was also determined by immunostaining and shown for each mouse. (C) C57BL/6 mice (n=2) were injected intraperitoneally with 200ug of Poly (I:C) and sacrificed at Day 2. Splenocytes were then stimulated for 4hours with plate bound NKp46 antibody. Intracellular staining for IFN-γ was performed. The proportion of IFN-γ+ NK cells among each maturation subsets (R1-R3) are shown for each individual. All data were analyzed using a paired non-parametric Wilcoxon test. p-values are shown on the graph. p-values are indicated on each graph.

**Figure S3:**
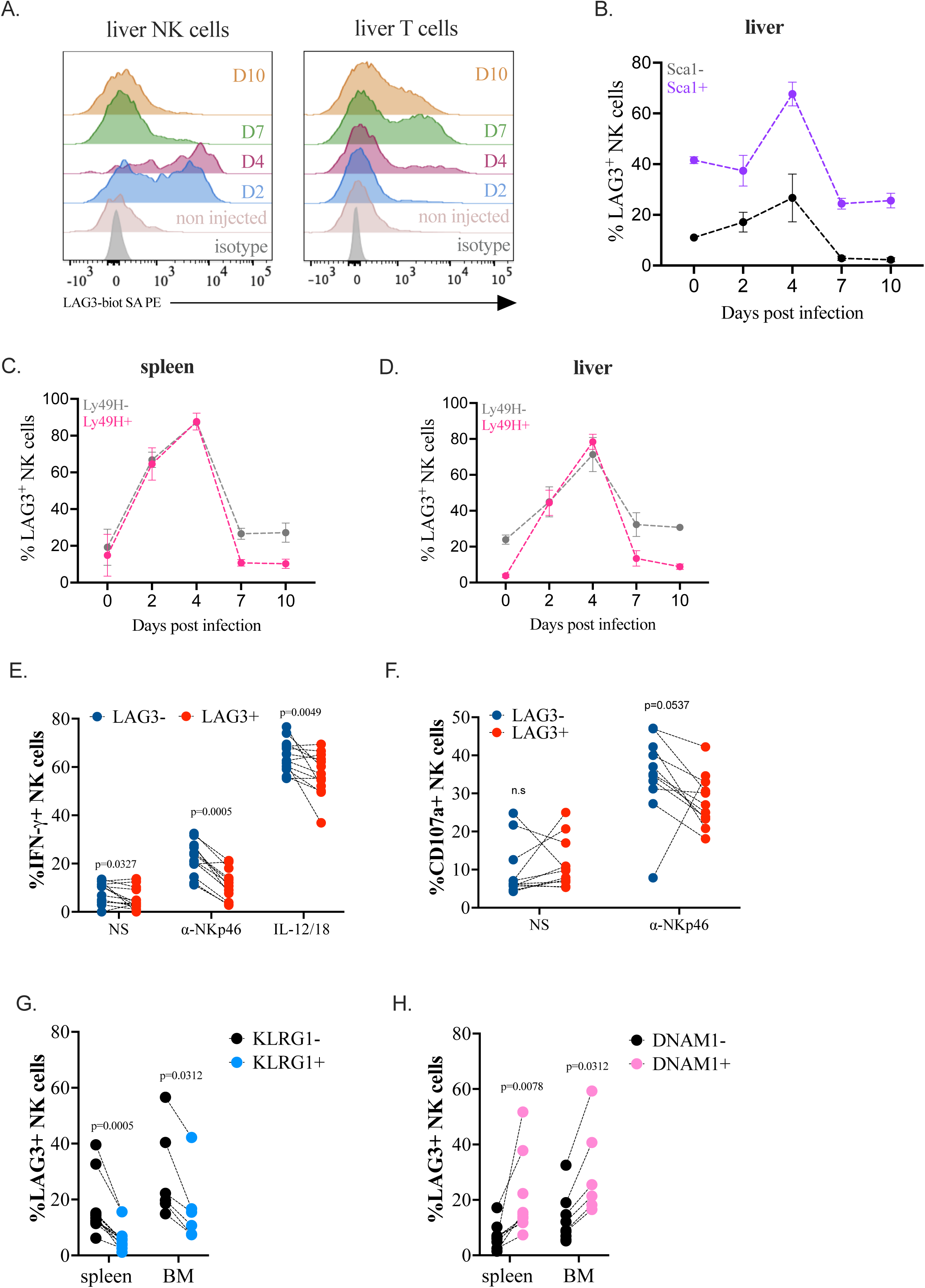
C57BL/6 mice (n=3 for each day analyzed) were infected with MCMV (15,000 PFU) and sacrificed in kinetic at D2, D4, D7 and D10. The expression of LAG3 on liver NK and T cells was determined by flow cytometry. Overlays of representative histograms are shown in NK cells (A, left panels) or in T cells (B, right panels). (B) The percentage of LAG3 among Sca1+ or Sca1-liver NK cells was determined by flow cytometry. Due to the small sample size, we did not calculate p-values. (C-D) The percentage of LAG3 among Ly49H+ or Ly49H-NK cells in the spleen (C) or liver (D) was determined by flow cytometry. Due to the small sample size, we did not calculate p-values. (E-F) Eµ-myc mice (n=11) were sacrificed at 12 weeks. Splenocytes were then stimulated for 4hours with plate bound NKp46 antibody or with IL-12+IL-18 (E) Intracellular staining for IFN-γ was performed. The proportion of IFN-γ + NK cells are shown for each individual. (F) the proportion of NK cells expressing CD107a was also determined by immunostaining and shown for each mouse. (G-H) Eµ-myc mice (n=9) were sacrificed at 12 weeks. The percentage of LAG3+ among KRLG1- and KLRG1+ NK cells (G) or DNAM1- and DNAM-1+ NK cells (H) in the spleen and bone marrow (BM) was determined by flow cytometry. All data were analyzed using a paired non-parametric Wilcoxon test. p-values are shown on the graph.

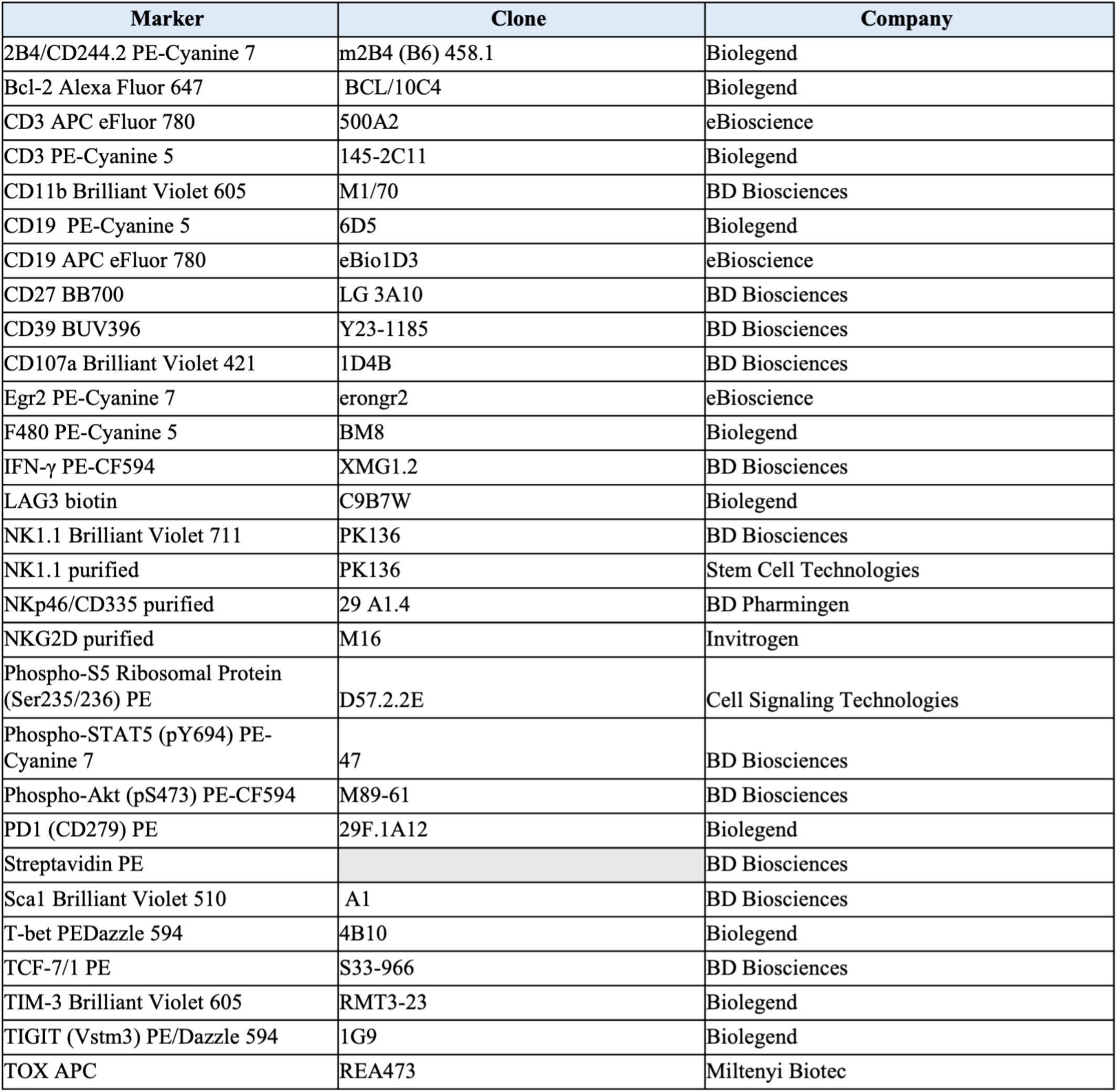

